# Sensory responsivity and its connection to sympathetic activation and deactivation: implications for stress and attention

**DOI:** 10.1101/2025.04.01.646564

**Authors:** Ellen Egeland Flø

## Abstract

This study introduces sensory responsivity (SR), which describes individual differences in sensory stimuli response strength and is hypothesised to affect stress responses in students and, in turn, their attention. To investigate this, a scale to assess SR was developed and linked to physiological responses. To this end, electrodermal activity (EDA) and sensory gating data was collected from a laboratory study with 100 students (12-21 y) and EDA data was collected in a classroom study with 35 students (17-18 y). In the lab study, sympathetic activation was generally lower for high SR groups, whilst in the classroom study, sympathetic activation was higher for high SR groups in line with the differential susceptibility theory. The high SR groups demonstrated higher sensory gating values, indicating a potential attentional benefit in low sensory input contexts. Thus, sensory responsivity moderates sympathetic activation based on the environmental sensory-intensiveness, which may impact attention and stress-related outcomes.

## 1. Introduction

Several groups are thought to be highly reactive to sensory stimuli, such as highly intelligent people (Beljan et al., 2006; Danthiir et al., 2000; Li et al., 1998; Helmbold et al., 2006), people with autism specter disorder (ASD) or attention deficit hyperactivity disorder (ADHD) (Dellapiazza et al., 2021; Scheerer et al., 2024), or people high in the personality trait of sensory processing sensitivity (Aron & Aron, 1997; Greven et al., 2019). There have been found extreme sensitivity to, e.g., taste, touch, and sound for the gifted (Beljan et al., 2006), where the ability to differentiate between similar sensory inputs, olfactory discrimination (Danthiir et al., 2000), tactile discrimination (Li et al., 1998), and temporal and pitch discrimination (Helmbold et al., 2006) have all been linked to intelligence.

Independent of intelligence, there have been identified connections between sensory issues and diagnoses such as ASD and ADHD (Dellapiazza et al., 2021; Scheerer et al., 2024). Descriptions of ASD individuals include issues with hypersensitivity, hyposensitivity, and sensory integration (Ben-Sasson et al., 2019; Schaaf et al., 2015). While self-reported atypical sensory processing is strongly linked to ASD, its physiological basis is unclear (Hazen et al., 2014), with links to behavioural and neural correlates being difficult to identify (Sapey□Triomphe et al., 2023). Similar results are found for ADHD, namely that sensory hyper- and hyposensitivity is common (Rani et al., 2023), persists into adulthood (Kamath et al., 2020), and is found independently of ASD symptoms (Bijlenga et al., 2017). More information on the underlying processes has been called for (Schulze et al., 2020).

A more general population with high reactivity to stimuli are those with elevated levels of sensory processing sensitivity (SPS) as defined by the top 20 % scoring of the HSP (highly sensitive person) scale (Aron & Aron, 1997; Greven et al., 2019). This scale includes items on positive or negative, emotional or cognitive responses to stimuli such as art, loud noises, smells, and fabrics (Greven et al., 2019). Examples of items are: “Do other people’s moods affect you?”, “Do you make a point to avoid violent movies and TV shows?”, and “Are you conscientious?”. SPS has been linked to brain activation patterns (Acevedo et al., 2018; Schaefer et al., 2022) but has been criticised for mainly describing emotional reactions to stimuli (Turjeman-Levi et al., 2022) and needing a more precise conceptualisation (Ershova et al., 2018; Hellwig & Roth, 2018).

Thus, in the current study, the concept of sensory responsivity, which describes individual differences in sensory stimuli response strength, is introduced. This includes the use of the sensory responsivity (SR) scale, which was developed to focus specifically on response strength to sensory stimuli from the visual, auditory, gustatory, olfactory, tactile, proprioceptive, and interoceptive sensory systems for the general population (see Appendix A for details). The results from the scale development and external validation showed that its psychometric properties were satisfactory, and two factors emerged, namely SR1, mainly describing exteroceptive items and SR2, mainly describing interoceptive items.

Through the external validation of the scale development process described in Appendix A, it was found that both SR scale factors correlated significantly with the three factors of the HPS scale (Kendall’s tau = 0.10 – 0.33), with the negative school engagement behaviour scale (Kendall’s tau = 0.17 for both factors) (Skinner et al., 2009), and with (Kendall’s tau = 0.074 – 0.091) suggesting that SR is a risk factor for school dropout.

Because there is evidence of individual differences in responses to sensory stimuli, but simultaneously a lack of conceptual clarity this study aims to introduce the concept of sensory responsivity and link it to activation of the sympathetic nervous system. To this end, the neurological and physiological measures of sensory gating and electrodermal activity are used, and implications for stress and attention are discussed.

Sensory gating is a neural process that filters out redundant stimuli from the environment, which is central for preventing sensory overload (Steiner & Tseng, 2016). It utilises the brain’s preconscious reaction to stimuli with its ability to reduce the response to a repeated auditory stimulus compared to the first (Freedman et al., 1987). Large gating values correspond to impaired sensory gaiting, for instance identified in children with sensory processing issues (Crasta et al., 2021; Davies et al., 2009). Another study linked reduced sensory gating to higher creative achievement and suggested the mechanism to be that the reduced gating helped people integrate ideas that are outside the attentional focus (Zabelina, et al., 2015), which is in line with research positively linking attention to sensory gating (Golubic et al., 2019; Jones et al., 2016; Wan et al., 2008). More generally, sensory gating is reduced at higher stress levels (Johnson & Adler, 1993; Xin et al., 2021), which makes it a befitting measure for the effects of sympathetic activation in the current study.

Electrodermal activity (EDA) sensors measure the conductivity of the skin. This conductivity is connected to activation of the sympathetic nervous system through the tonic and phasic firing of the eccrine sweat glands, which emit more sweat when aroused (i.e., the sympathetic nervous system is activated) (Benedek & Kaernbach, 2010; Boucsein, 2012; Ellaway et al., 2010). Furthermore, sympathetic activation governs processes that non-consciously regulate the mobilisation of the human body for action (Dawson et al., 2000), and has been tied to attention (Villanueva et al., 2018), psychological stress (Yang et al., 2015), and physiological stress (Godoy et al., 2018).

When sympathetic activation is unnecessarily high, particularly over time, there are negative associations with these sustained stress levels. For example, perceived stress is associated with sympathetic activation and identified as a risk factor for coronary artery disease (Yang et al., 2015), sympathetic activation is also associated with the severity of coronary artery disease (Boieriu et al., 2024), and heightened sympathetic activation is a critical contributor to the link between stress and cardiovascular disease (Hering et al., 2015). Not only is the cardiovascular system affected by stress, but it has been shown that the sympathetic nervous system can modulate cutaneous immune responses and that the development and progression of atopic dermatitis and psoriasis are impacted by stress (Hall et al., 2012). More generally, stress is associated with disease onset and exacerbation in other autoimmune conditions such as rheumatoid arthritis, systemic lupus erythematosus, inflammatory bowel disease, multiple sclerosis, and Graves’ disease (Sharif et al., 2018).

That is, if sensory responsivity mediates the individual’s sympathetic activation level, it could be relevant to stress-related research and the medical and biological responses to stress (e.g., cardiovascular health and autoimmune conditions), as well as research on attention. To introduce sensory responsivity and understand its connection to sympathetic activity, stress, and attention, the following hypotheses are investigated.

### Hypotheses

#### H1

The highest SR groups have lower levels of sensory gating.

#### H2

The highest SR groups have higher levels of sympathetic activation, and the difference depends on the sensory intensity of the environment.

These hypotheses were addressed through a lab study which investigated the connection between sensory responsivity and sensory gating, as well as sensory responsivity and sympathetic activation in two relaxation and two concentration tasks. Because the lab experiment took place in a very quiet lab, where sensory input was generally low, a classroom study was also conducted to investigate sympathetic activation for learning tasks in a naturalistic setting with higher levels of sensory input, in order to fully investigate hypothesis two.

## 2. Method

### 2.1 Participants and procedure

The study was approved by the south-east Regional Ethics Committee (ref. removed for blinded review) and the Norwegian Centre for Research Data (ref. removed for blinded review) and students (and their guardians if they were below 16 years) provided written informed consent for participation. We gave each participant a unique and anonymous ID. For the students not consenting to video data collection in the classroom study, they were placed outside the stationary camera’s recording angle.

#### 2.1.1 Lab study

The data was collected by three research assistants at a cognitive lab with 100 participants, where 59 were female, 39 were male, and two preferred not to disclose. The participants were aged 12 – 21 years (M = 16.5, SD = 1.7) and were recruited through their schools, which sent out information about the project. This included information that students with diagnoses of ADHD, ASD, ME/CFS, dyslexia, any learning disabilities, depression, anxiety or similar psychiatric diagnoses were not eligible for participation. When scheduling the data collection, the exclusion criteria were included in the communication, and when collecting the survey data, students were required to confirm that they did not have any of these diagnoses or learning disabilities. The data collection lasted about 50 minutes, and the participant was seated in a chair with an approximate distance of 60 cm between the computer monitor and their eyes. See Figure 1 for the lab setup.

**Figure 1.**
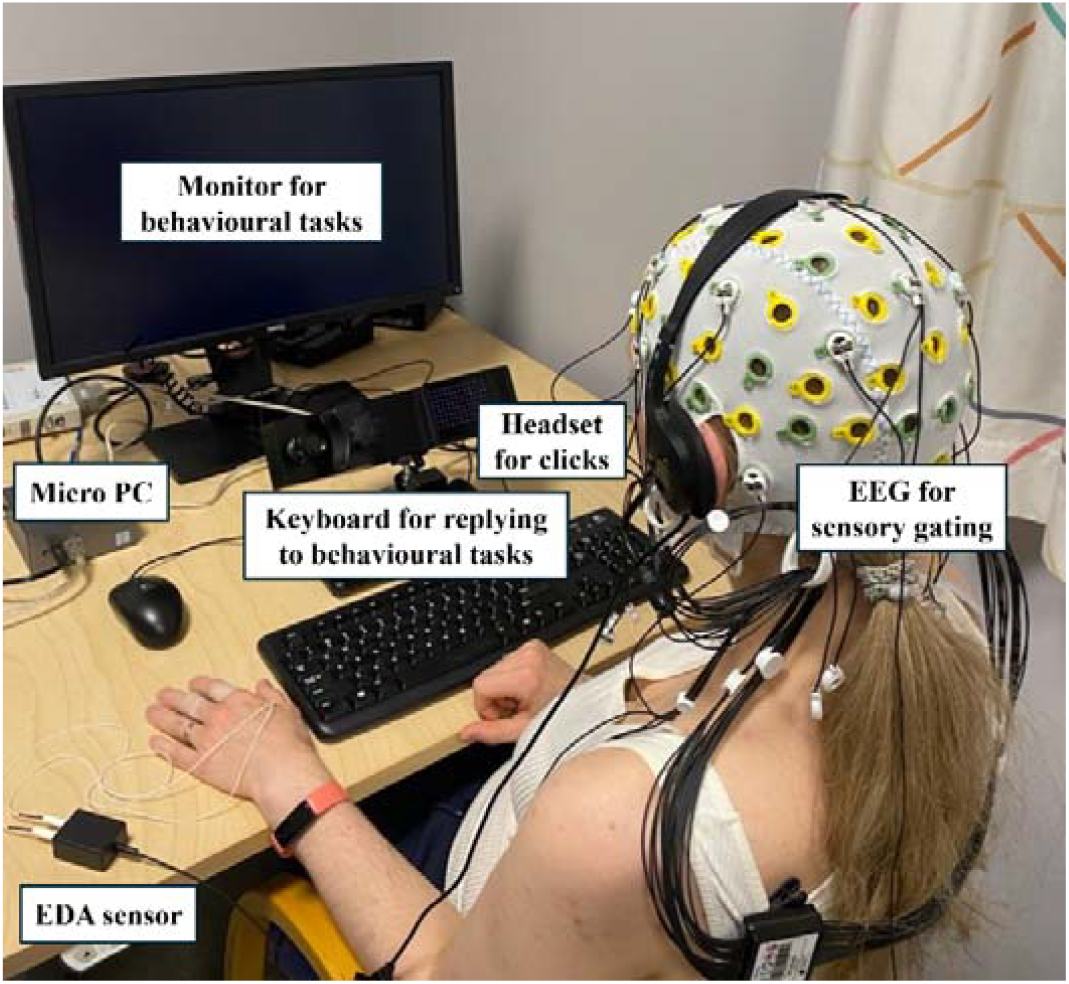
The lab setup (with the author as participant) included a monitor at a 60 cm distance from the participant. The EDA sensors were fastened to the fingers of the non-dominant hand, but both hands were required to press keys on the keyboard to answer some of the behavioural tasks. The lab had no windows, and light and temperature levels were kept constant.

The participants performed four tasks, namely, watching a one-minute relaxing nature video, a sensory gating task, watching a six-minute learning video in biology, and performing an N2-back test. Behavioural results from the relaxing video, learning video and N2-back task will be reported in other works.

The sensory gating paradigm consisted of passively listening to two auditory clicks per trial, each a 1 ms square wave with an intensity of 90 dB. The trials were randomly given at 8 ± 2 seconds with 500 ms between the paired clicks. The participants were instructed to sit upright and alert, but with closed eyes to avoid blinking artefacts. The sensory gating task consisted of 99 trials, which lasted approximately 14 minutes.

#### 2.1.2 Classroom study

35 students (19 female) aged 17-18 years from two upper secondary school physics classes participated in an instructional activity intervention where one class was first given activities with less intense sensory stimuli, and the next time, they were given the same instructional activities only with more intense sensory stimuli. The other class received the same intervention, only in the reverse order. The order of the themes the students worked with was kept the same for both groups in such a way that both groups had theme one the first time and theme two the second time (see Figure 2). Both lessons consisted of a lecture with class interaction (25 min -students could take notes on laptop vs. by hand), worked examples at the whiteboard (10 min - students could not take notes vs. taking notes by hand), working with tasks with four different difficulty levels (20 min - using laptop with digital book vs. by hand with printed book), a quiz (10 min - Kahoot! vs. sitting on their desks and showing answers with their arms), and a round of the Alias game with appropriate content (10 min - explaining concepts orally vs. explaining orally and drawing).

**Figure 2.**
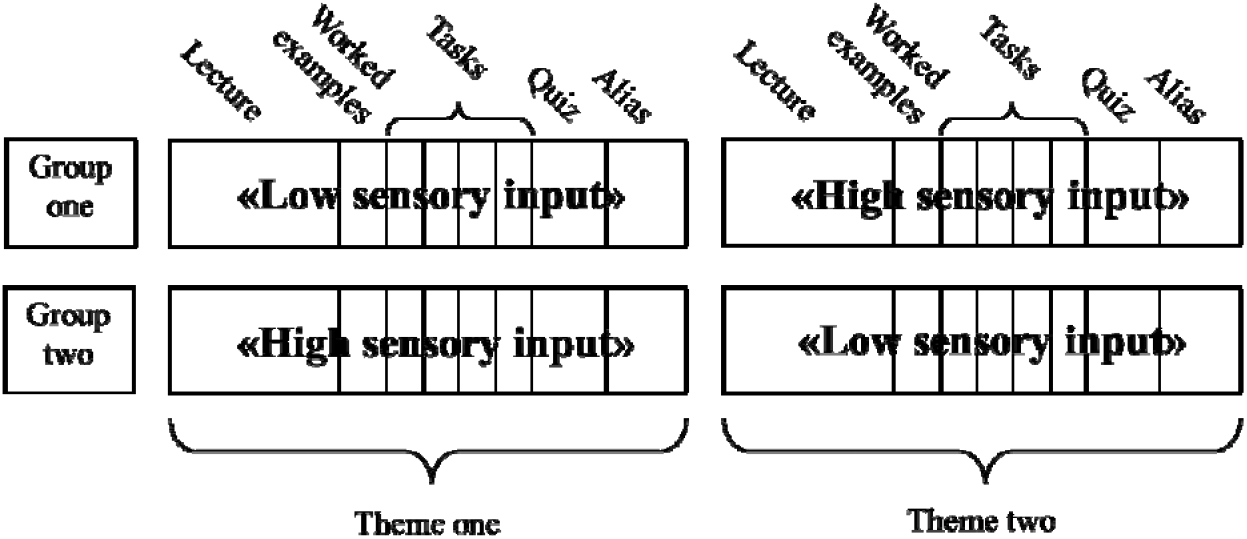
An overview of the experimental design where each of the four lessons had the same learning activities, i.e., lecture with students asking questions and discussing with the teacher, worked examples on the whiteboard with input from students, physics tasks of four different difficulty levels where students could cooperate if they wanted to, a quiz where the answers were not collected nor graded, and a game of alias in groups of 3-4 students. These activities were either done in a low sensory input condition with notetaking on laptops during the lecture, not taking notes during worked examples (but students would receive the stepwise solutions after class), using laptop and digital book when solving tasks, a Kahoot-quiz, and explaining the concepts orally in a game of “Alias”. Otherwise these activities were done in a high sensory input condition with notetaking by hand during the lecture, notetaking by hand during worked examples, using printed book and solving tasks by hand (could use calculator when needed), a quiz where students sat on their desks and help up their arms and hands to show their answer, and explaining the concepts both orally and by drawing when playing “Alias”.

### 2.2 Instruments

The SR scale demonstrated robust internal validity, and external validity through positive correlations with the HSP (highly sensitive person) scale (Aron & Aron, 1997), the Negative School Engagement Behaviour scale (Skinner et al., 2009), and self-reported school absence level, which are described in greater detail in Appendix A, page 8. Further details on the sample (N = 535, 9-16 years) and methods (multidimensional item response theory) used for development of the SR scale are described in Appendix A.

The spatial reasoning instrument (SRI) (Ramful et al., 2017) assesses spatial reasoning abilities in middle school students. It has previously been validated and used with Norwegian youth up to 16 years without reaching ceiling effects (Flø & Smedsrud, 2025). The SRI consists of 30 multiple-choice items measuring mental rotation, spatial visualisation, spatial orientation, and spatial relations (Ramful et al., 2017). It demonstrates robust validity and reliability (Ramful et al., 2017).

### 2.3 Data collection and processing

#### 2.3.1 EEG (sensory gating)

Three research assistants collected EEG data with a Brainproducts ActiChamp Plus system at 5000 Hz using the standard 64 channel actiCAP snap electrode cap with 14 electrodes. The electrodes were placed at FP1, FZ, FP2, F3, F4, F7, F8, C3, CZ, C4, P3, PZ, P4, and earlobes based on the 10/20 system (Zabelina et al., 2015). The ground electrode was fixed between the FP1 and FP2 electrodes. The impedance for all electrodes was kept at less than 5 kΩ. No online filtering was performed, whereas online reference was one ear lobe electrode, and offline reference was the mean of both ear lobe electrodes.

The BrainVision Analyzer 2.2 software (Brain Products, Gilching, Germany) was used for the evoked response potential (ERP) analysis. The individual EEG data was re-referenced offline to the mean of the two earlobe electrodes, which was suitable due to the midline focus of the ERP analysis (Luck, 2014), and in line with previous methodology (Chang et al., 2012; Olincy et al., 2010; Zabelina et al., 2015). Then, the data was filtered with a zero-phase shift Butterworth filter with a low cutoff of 1 Hz, high cutoff at 70 Hz, and a notch filter at 50 Hz to eliminate noise from the power grid.

Trials with artifacts (i.e., amplitude values beyond -75 to 75 µV) were visually inspected and removed from the analysis. After artifact rejection, EEG data was segmented into event-locked epochs for the first and second click, with epochs extending from 100 ms before to 200 ms after the stimulus. Following recommendations from previous literature (Gumenyuk et al., 2013; Lijffijt et al., 2009), non-rejected segments underwent baseline correction using the pre-stimulus interval (i.e., -100 to 0 ms). These corrected segments were averaged to generate ERP waveforms for each click.

Following established protocols (Chang et al., 2012; Kurthen et al., 2007; Miller et al., 2018; Olincy et al., 2010), P50 amplitudes were measured at Cz using the trough-to-peak method. This method involves identifying the P50 response as the most positive peak, in µV, of the ERP waveform within the 35–75 ms window following the first click onset. The amplitude was calculated by subtracting the amplitude of the most negative trough, which occurs immediately before the P50 peak within the same time window, from the peak amplitude. For the second click, the P50 response was consistently found within 10 ms of the peak response to the first click, and its amplitude was calculated using the same method.

Data integrity was ensured by inspecting the EEG topography for the relevant time frame and the phase locking factor (Chang et al., 2012) for the Cz channel, in addition to visually inspecting all waveforms. This was followed by averaging across trials, where the P50 peak was identifiable for both Click 1 and Click 2 in all participants. The final P50 ratio for each participant was calculated by dividing the trough-to-peak amplitude of the averaged second click by that of the averaged first click (S2/S1) (Chang et al., 2012; Olincy et al., 2010).

#### 2.3.2 EDA

In the lab study, EDA data was collected with a BP-MB-30 GSR module, including electrodes and gel at 5000 Hz through the Brainproducts ActiChamp Plus system. Electrodes were taped to the second and third proximal phalanges of the non-dominant hand of the participants with medical tape. In the classroom study, one research assistant and the author measured EDA with the validated BITalino EDA sensor (Batista et al., 2019) at 10 Hz with two solid-gel self-adhesive disposable Ag/AgCl electrodes. The temperature was kept at 22 °C for both studies to prevent thermoregulatory interference with the EDA measurement (Boucsein, 2012).

MATLAB-based software Ledalab (version 3.4.9; http://www.ledalab.de/) was used for EDA signal processing, with no preprocessing, as is the default of the Ledalab software (Benedek & Kaernbach, 2010). Because EDA was collected at 5000 Hz in the lab study, it was down-sampled to 10 Hz to speed up the analysis, whereas in the classroom study, EDA data was collected at 10 Hz and needed no down-sampling. To decompose the EDA signal into the phasic and tonic components, the recommended continuous decomposition analysis (CDA) was applied (Benedek & Kaernbach, 2010).

### 2.5 Statistical analysis

R version 4.3.1 (R Core Team, 2023) with R studio version 2023.12.1 was used for the statistical analysis. The data was assessed for normality using the Shapiro-Wilk test. To assess whether sensory responsivity predicted sensory gating or EDA levels, a linear mixed-effects model was estimated per outcome type in the lab study, and one model for the classroom study. Non-significant covariates were eliminated, and the resulting reduced model was compared to the original model with a likelihood ratio test. Afterwards, post-hoc tests with marginal means were conducted with Bonferroni-Holm corrections applied. For all group comparisons, a median split was performed for each SR scale factor (SR1 and SR2).

All regression models were checked for multicollinearity with the variance inflation factor (VIF), linearity with scatterplots and residual plots, homoscedasticity with the non-constant variance test, and normality through visually inspecting the Q-Q plots and the Shapiro-Wilk test.

For sensory gating, initial regression diagnostics revealed systematic violations of normality, heteroscedasticity, and influential observations. Transformations (i.e., log, Box-Cox) and heteroscedasticity-consistent standard errors failed to alleviate these issues. Robust regression (i.e., lmrob) and bias-corrected bootstrap confidence intervals produced unstable coefficient estimates and asymmetric intervals, indicating that the sampling distribution of the mean outcome was non-normal. Accordingly, the Mann-WhitneyLU test was adopted as the primary inferential procedure for the sensory gating analysis as it does not rely on the assumptions violated in the parametric models. The corresponding effect size, the rank biserial correlation, r (Tomczak & Tomczak, 2014), was calculated.

## 3. Results

The results are structured according to the two research hypotheses, by first focusing on whether the highest SR groups have lower levels of sensory gating in section 3.1, then focusing on whether the highest SR groups have higher levels of sympathetic activation in section 3.2 for the lab study in a quiet environment and in section 3.3 for the more sensory intensive environment in the classroom study.

### 3.1 Sensory gating – lab study

The spatial distribution of neural activity across the scalp was first analysed to examine the neural correlates of sensory gating in the P50 response. The EEG topography maps for the P50 response timeline are presented in Figure 3, with the Cz electrode positioned at the centre of the head as per standard double-click auditory ERP methodology (Korzyukov et al., 2007; Patterson et al., 2008; Spooner et al., 2020). These topographical representations were used to ensure signal stability throughout the analysis period. Following this initial verification of spatial distribution, the sensory gating effects were quantified by comparing the amplitude of responses to the first (S1) and second (S2) auditory clicks across different sensory responsivity groups.

**Figure 3.**
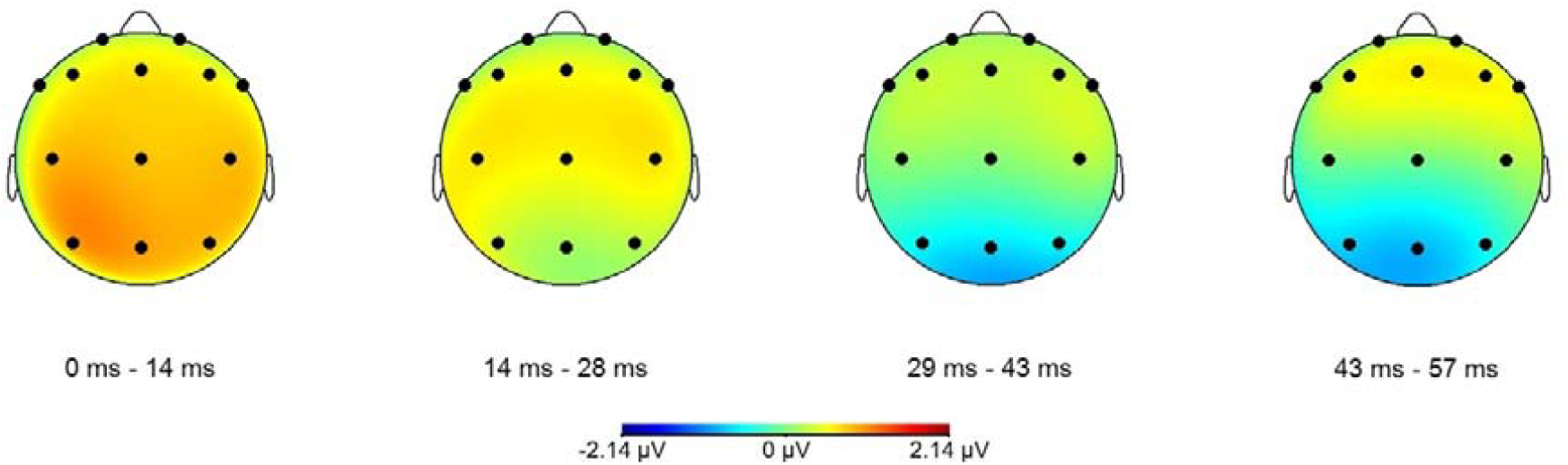
EEG topography maps for the P50 response timeline. The maps illustrate the spatial distribution of neural activity during the P50 response, with the Cz electrode positioned at the centre of the head as per standard double-click auditory ERP methodology. These topographical representations were utilised to ensure signal stability throughout the analysis period.

Since the P50 signal is naturally small and mixes with background oscillations, a phase locking factor (PLF) was performed to verify if evoked activity is systematically produced after stimulation at the Cz electrode site. This revealed phase locking (event-related) of about 40 Hz at about 50 ms post-stimulus, for both the first and the second click, as shown in Figure 4. Since the P50 shows frequencies of about 40 Hz (Chang et al., 2012), this observation supports that the P50 component is being evoked after the stimuli markers.

**Figure 4.**
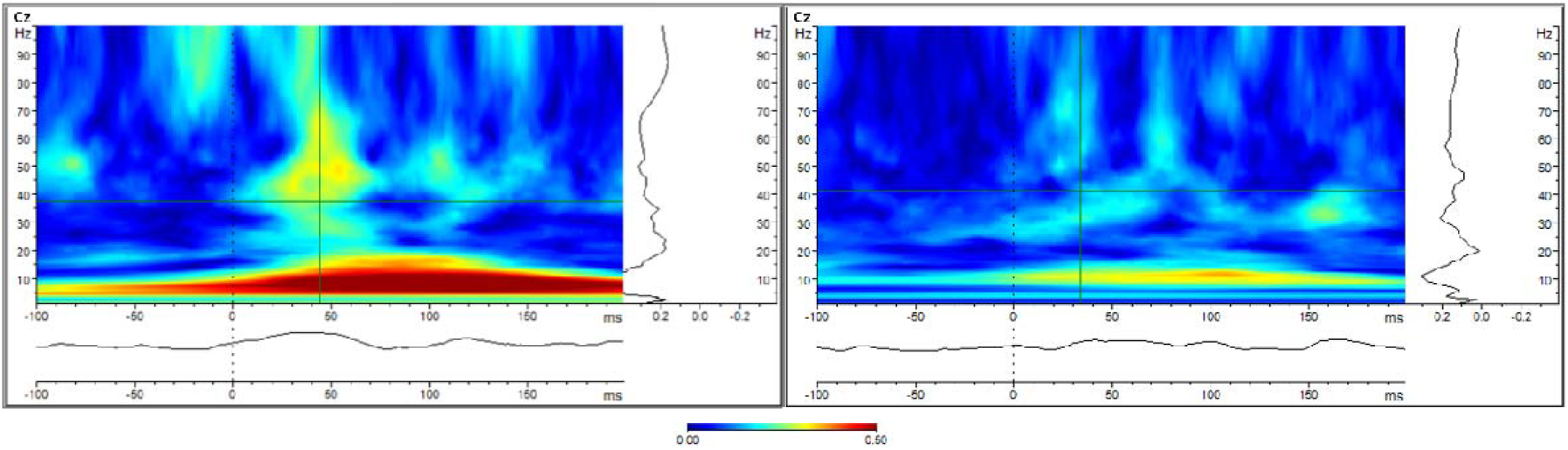
Grand averages of the phase-locking factors obtained from ERPs at Cz. The vertical dashed lines at 0 ms mark onset of the stimuli (click 1 on the left panel and click 2 on the right panel).

The sensory gating effect is characterised by a significant reduction in amplitude of the P50 response to the second click (S2) compared to the first click (S1), reflecting the brain’s ability to filter out repetitive stimuli (Freedman et al., 1987). Analysis of the grand average ERP waveforms for the P50 response revealed that both high and low sensory responsivity groups demonstrated effective sensory gating, evidenced by the typical attenuation of the S2 response relative to S1, as seen in Figure 5.

**Figure 5.**
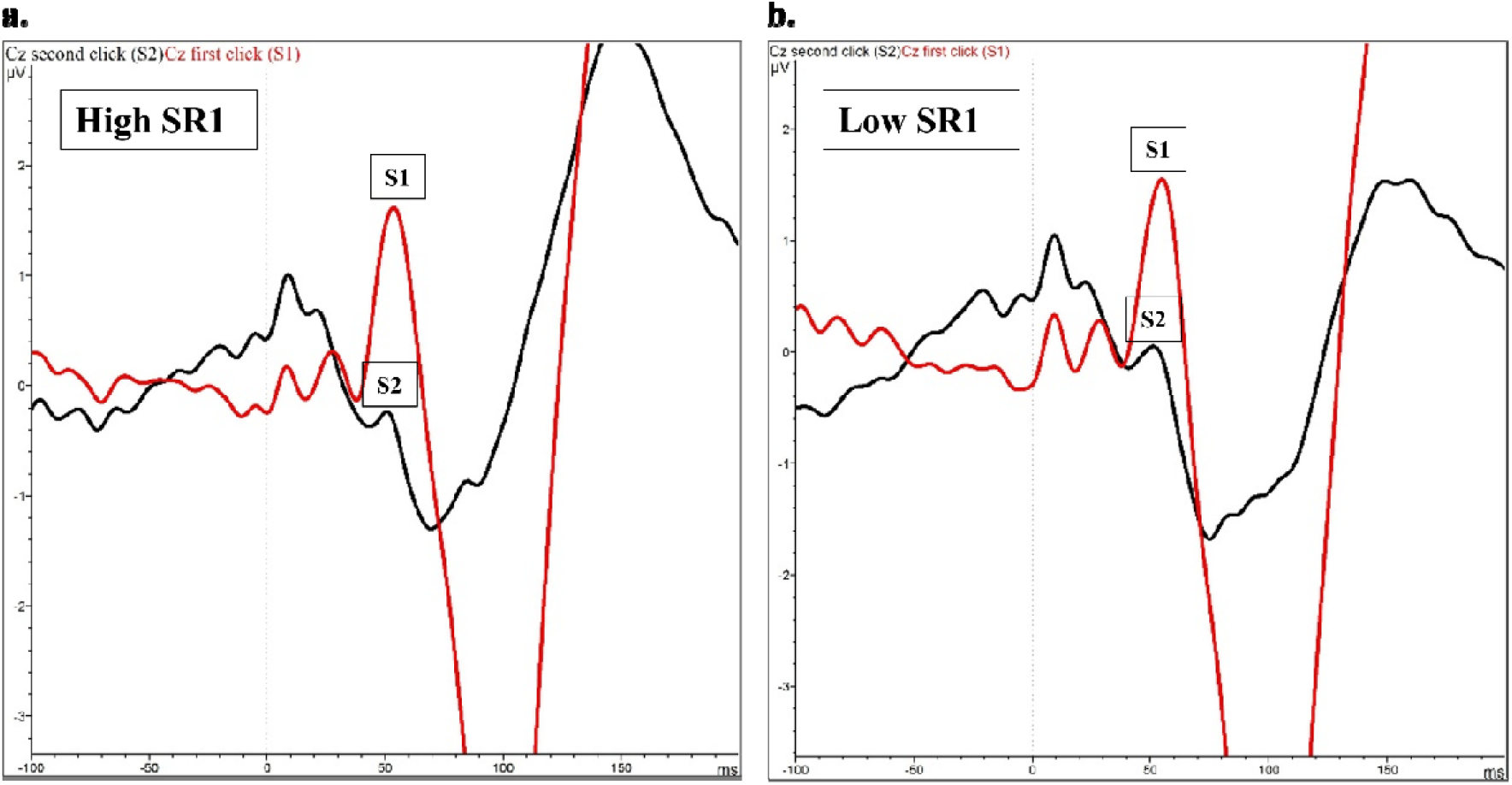
Grand averages of the ERPs at Cz. Vertical dashed lines at 0 ms mark onset of both clicks. The P50 sensory gating is calculated as the ratio of the P50 peak to trough difference of click 2 to the P50 peak to trough difference of click 1 (S2/S1). **Panel a**. The P50 response of the first click (S1) and the second click (S2) for the high SR1 group. **Panel b**. The P50 response of first click (S1) and the second click (S2) for the low SR1 group.

The results in Figure 6 show that the amplitude of the P50 response to the first auditory click (S1) in the paired-click paradigm was higher for the high SR1 group compared to the low SR1 group, which was also the case for the high and low SR2 groups, respectively. Statistical analyses using Mann-Whitney U tests (due to non-identical groups and non-normal distributions) with a Bonferroni-Holm correction confirmed these group differences for both SR1 groups, V = 113,050, p < .001, and SR2 groups, V = 113,050, p < .001. The effect sizes were small for both comparisons, with r = .134 (95% CI [.006, .33]) for the SR1 comparison and r = .235 (95% CI [.05, .43]) for the SR2 comparison. For the second auditory click (S2), the same pattern of results was observed, with significant differences between high and low SR1 groups, V = 113,050, p < .001, r = .128 (95% CI [.005, .31]), and between high and low SR2 groups, V = 113,050, p < .001, r = .238 (95% CI [.05, .43]).

**Figure 6.**
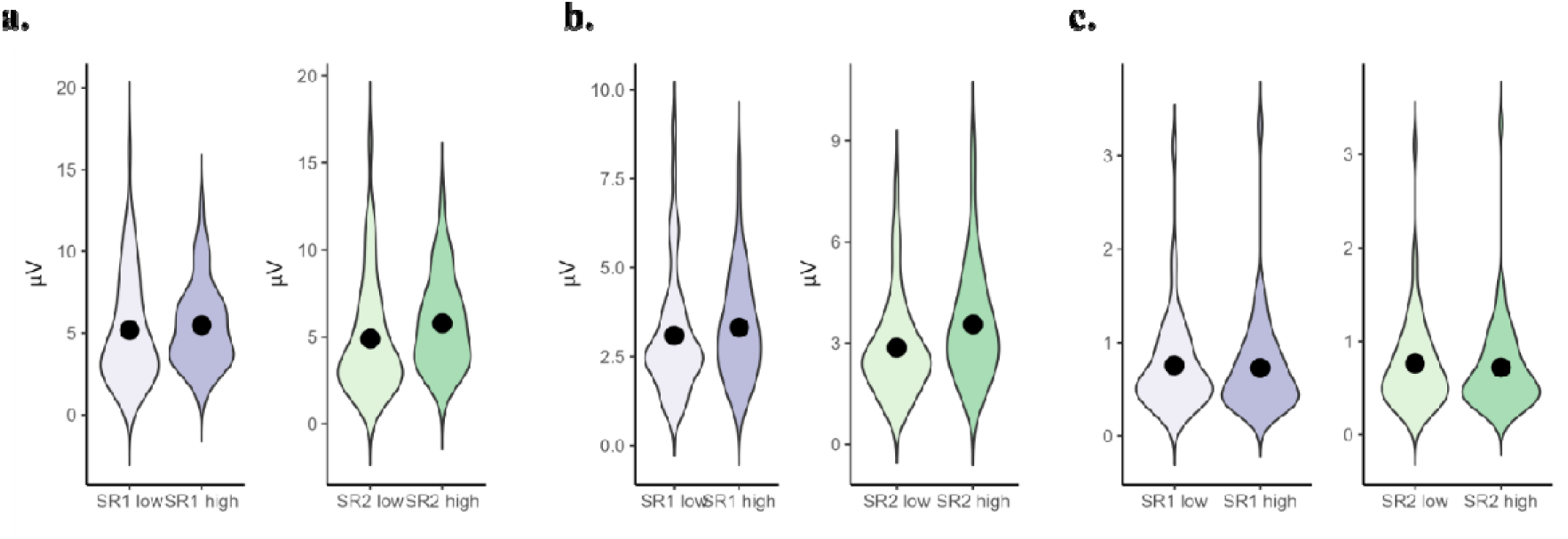
The preconscious response to auditory stimuli. **Panel a**. The P50 response of the first click (S1) for the SR1 and SR2 dichotomised groups. **Panel b**. The P50 response of the second click (S2) for the SR1 and SR2 dichotomised groups. **Panel c**. The sensory gating for the SR1 and SR2 dichotomised groups. All group differences were statistically significant.

Finally, the sensory gating (S2/S1) was, however, lowest for both high SR groups compared to the corresponding low SR groups, meaning that participants with higher SR demonstrated better sensory gating, i.e., that S2 is more dampened relative to S1, as shown in the test for SR1, revealed a significant difference between the groups, V = 113,050, p < .001, with a small effect size, r = .044, 95% CI [.003, .26], and the test for SR2, revealed a significant difference between the groups, V = 113,050, p < .001, with a small effect size, r = .050, 95% CI [.004, .23].

### 3.2 Sympathetic activation – lab study

Sympathetic activation manifests as tonic (sustained) and phasic (transient) components that serve as physiological markers of autonomic arousal (Boucsein, 2012). Analysis of the EDA waveforms revealed distinct patterns of sympathetic activation between high and low sensory responsivity groups, with mainly tonic EDA components showing differential responses across the groups in a low sensory intensity context, as illustrated in Figure 7.

**Figure 7.**
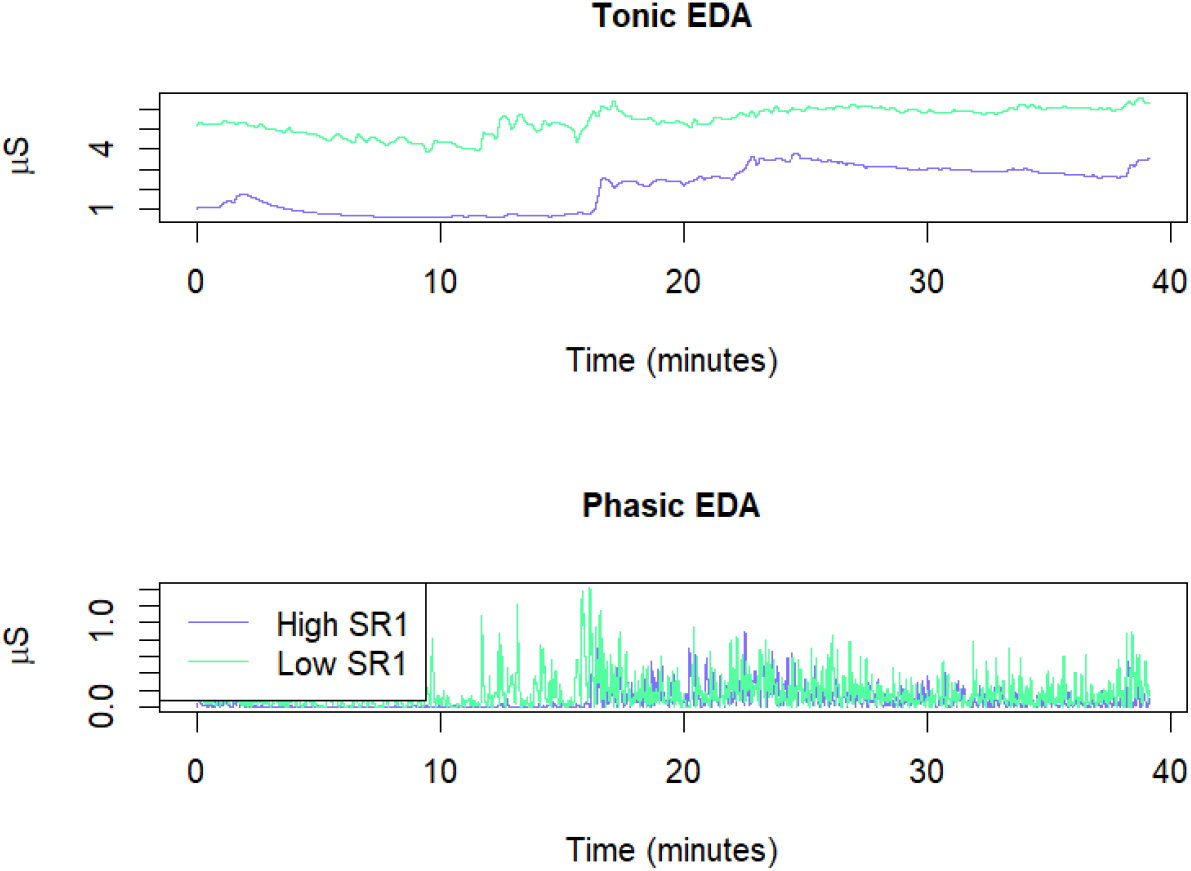
An example of tonic and phasic EDA responses for one high sensory responsivity (SR1) student versus one low sensory responsivity (SR1) student in the lab study.

To assess whether sensory responsivity predicted tonic or phasic EDA levels, a linear mixed-effects model was estimated as the two EDA measures are not independent, with EDA as outcome. This model controlled for spatial ability, age, gender, EDA type (tonic or phasic), activity (learning video, N2-back test, relaxing video, sensory gating) and an interaction between the EDA type and activity, and another interaction between the EDA type and the SR group factor (median-split into high and low), and investigated main effects from the SR group factor (two different models, as these variables demonstrated high multicollinearity (r = .903).

Spatial ability, gender, and age were not significant predictors of EDA. A likelihood ratio test comparing the full mixed model and a reduced model without spatial ability and age revealed that the full model (χ^2^(3) = 3.08, p = .214 for SR1 and χ^2^(3) = 3.28, p = .193 for SR2) did not provide a significantly better fit to the data than the reduced model without the additional predictors. This comparison therefore justified dropping the spatial ability and age variables and proceeding with the simpler models.

The simple mixed-effects models demonstrated significant contributions to the EDA outcome from all variables. For the SR1 model, the conditional R^2^ (which includes both fixed and random effects) was .807, while the marginal R^2^ (which includes only fixed effects) was .678. Similarly, for the SR2 model, the conditional R^2^ was .808, while the marginal R^2^ was .680. Model diagnostics indicated no violations of linearity, homoscedasticity, or normality of residuals. The variance inflation factors were all below 2, suggesting that multicollinearity did not bias the estimates.

The linear mixed effects model for SR1 revealed significant main effects of SR1 (b = -.754, SE = .239, p = .002), EDA type (b = -4.830, SE = .198, p < .001), and condition (sensory gating: b = -.570, SE = .177, p = .001; N2-back: b = .519, SE = .177, p = .003). The model also identified significant interaction effects between SR1 and EDA type (b = .750, SE = .177, p < .001), and between EDA type and activity (sensory gating: b = .542, SE = .250, p = .030; N2-back: b = -.504, SE = .250, p = .043).

The linear mixed effects model for SR2 revealed significant main effects of SR2 (b = -.825, SE = .238, p < .001), EDA type (b = -4.865, SE = .197, p < .001), and activity (sensory gating: b = -.570, SE = .176, p = .001; N2-back: b = .519, SE = .176, p = .003). The model also identified significant interaction effects between SR2 and EDA type (b = .819, SE = .176, p < .001), and between EDA type and activity (sensory gating: b = .542, SE = .249, p = .030; N2-back: b = -.504, SE = .249, p = .044).

Post-hoc pairwise comparisons with Bonferroni-Holm correction revealed that the difference between SR1 and SR2 groups was significant only for tonic EDA across all conditions. Specifically, for tonic EDA, the low SR1 group showed significantly higher EDA responses than the high SR1 groups (estimate = .754, SE = .242, t(130) = 3.12, p = .002) for the relaxing video, sensory gating, learning video, and N2-back task. In contrast, no significant differences were found between SR1 groups for phasic EDA in any of the conditions (estimate = .004, SE = .242, t(130) = .02, p = .988 for all conditions). Similarly, for tonic EDA, the low SR2 group showed significantly higher EDA responses than the high SR2 group (estimate = .825, SE = .241, t(130) = 3.43, p < .001) for the relaxing video, sensory gating, learning video, and N2-back task. In contrast, no significant differences were found between SR2 groups for phasic EDA in any of the conditions (estimate = 0.006, SE = .241, t(130) = .03, p = .980 for all conditions). These results indicate that both the SR1 and SR2 group differences are specific to tonic EDA responses and are consistent across all experimental activities, as shown in Figures 8 and 9.

**Figure 8.**
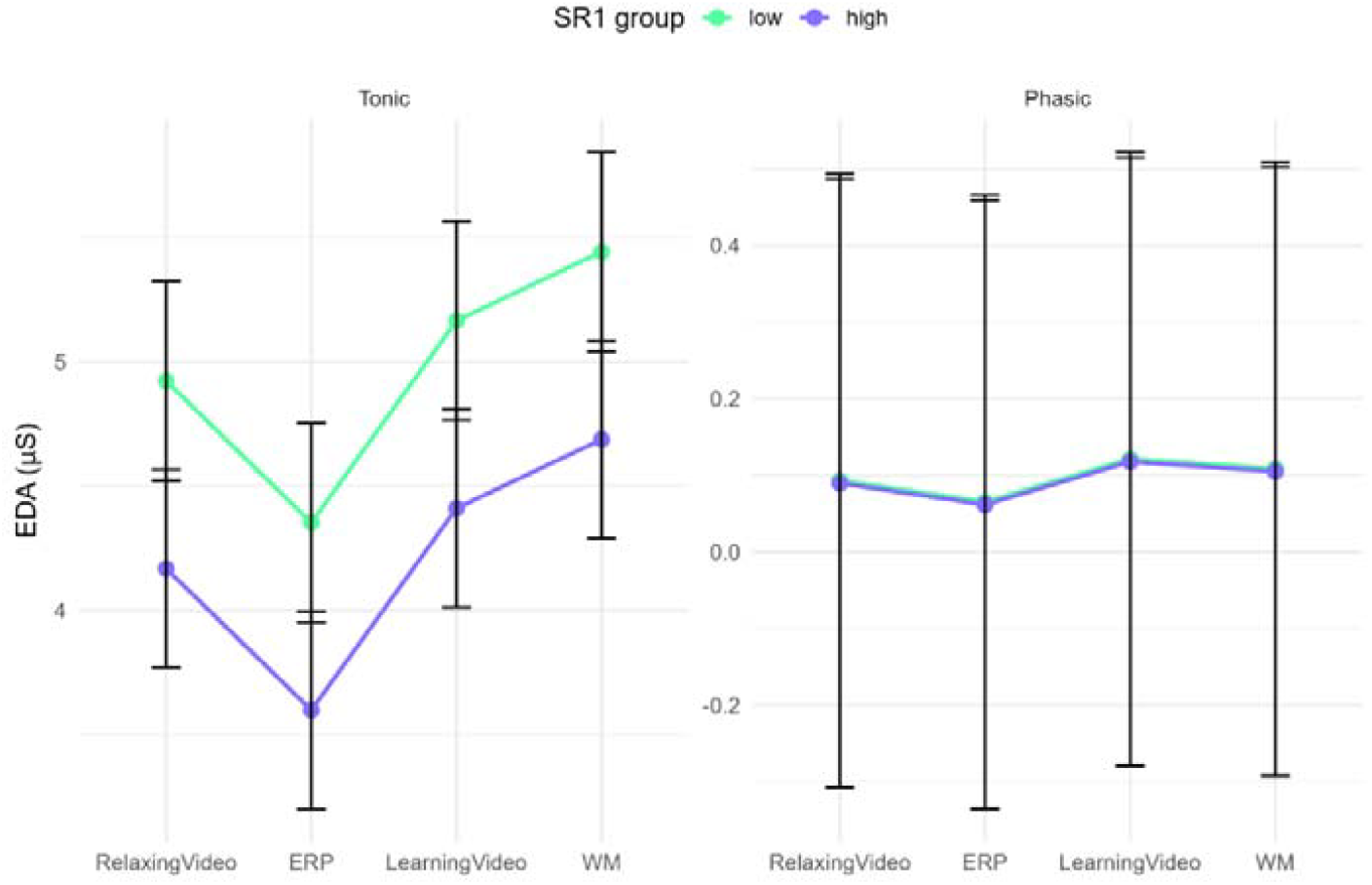
Tonic and phasic EDA response in the lab study across the four activities for the SR1 median split groups.

**Figure 9.**
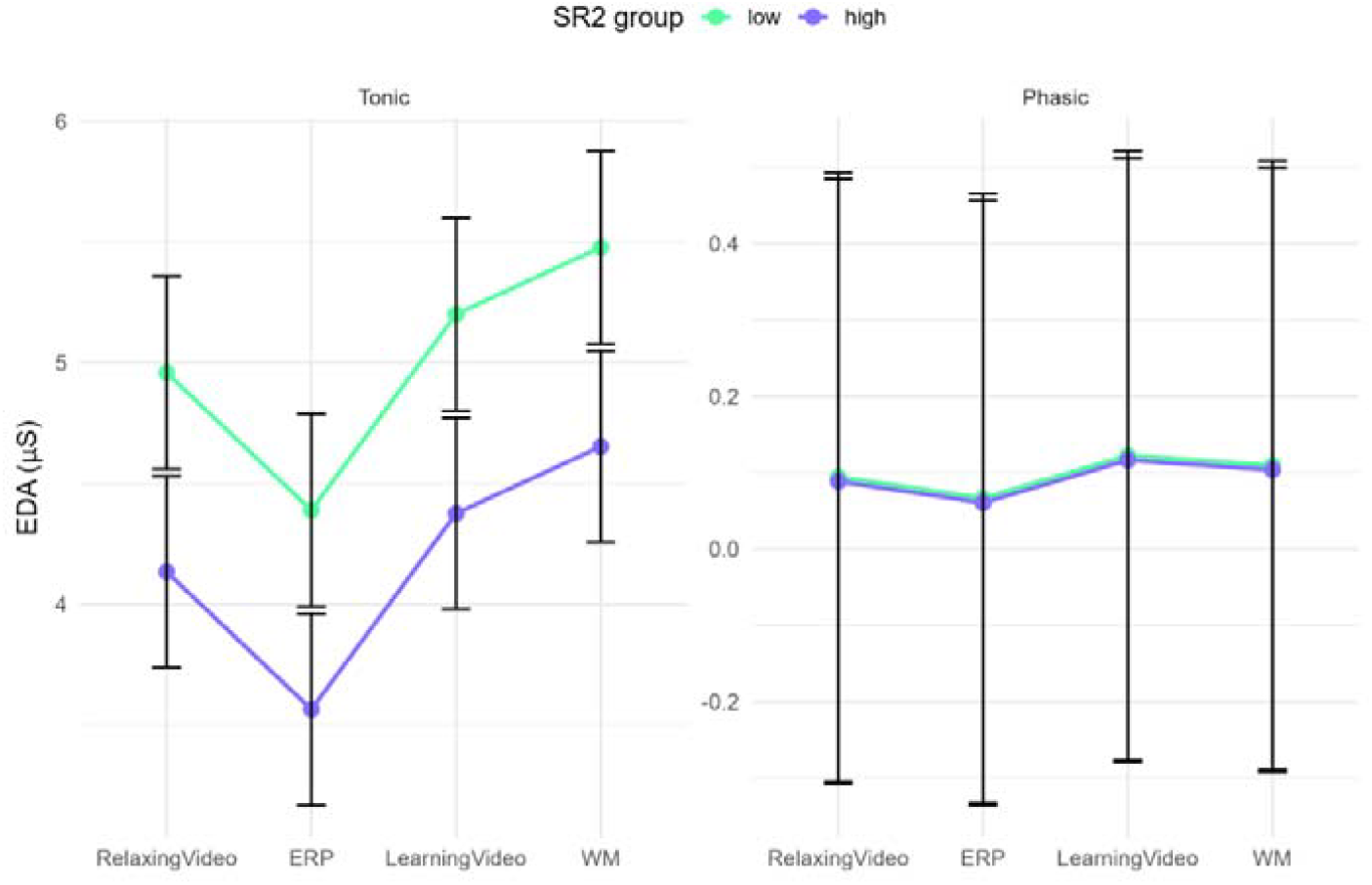
Tonic and phasic EDA response in the lab study across the four activities for the SR2 median split groups.

### 3.3 Sympathetic activation – classroom study

In the classroom study, where sensory intensity was higher than in the lab study, analysis of the EDA waveforms also revealed patterns of sympathetic activation between high and low sensory responsivity groups, with mainly tonic EDA components showing differential responses across the groups, as illustrated in Figure 10.

**Figure 10.**
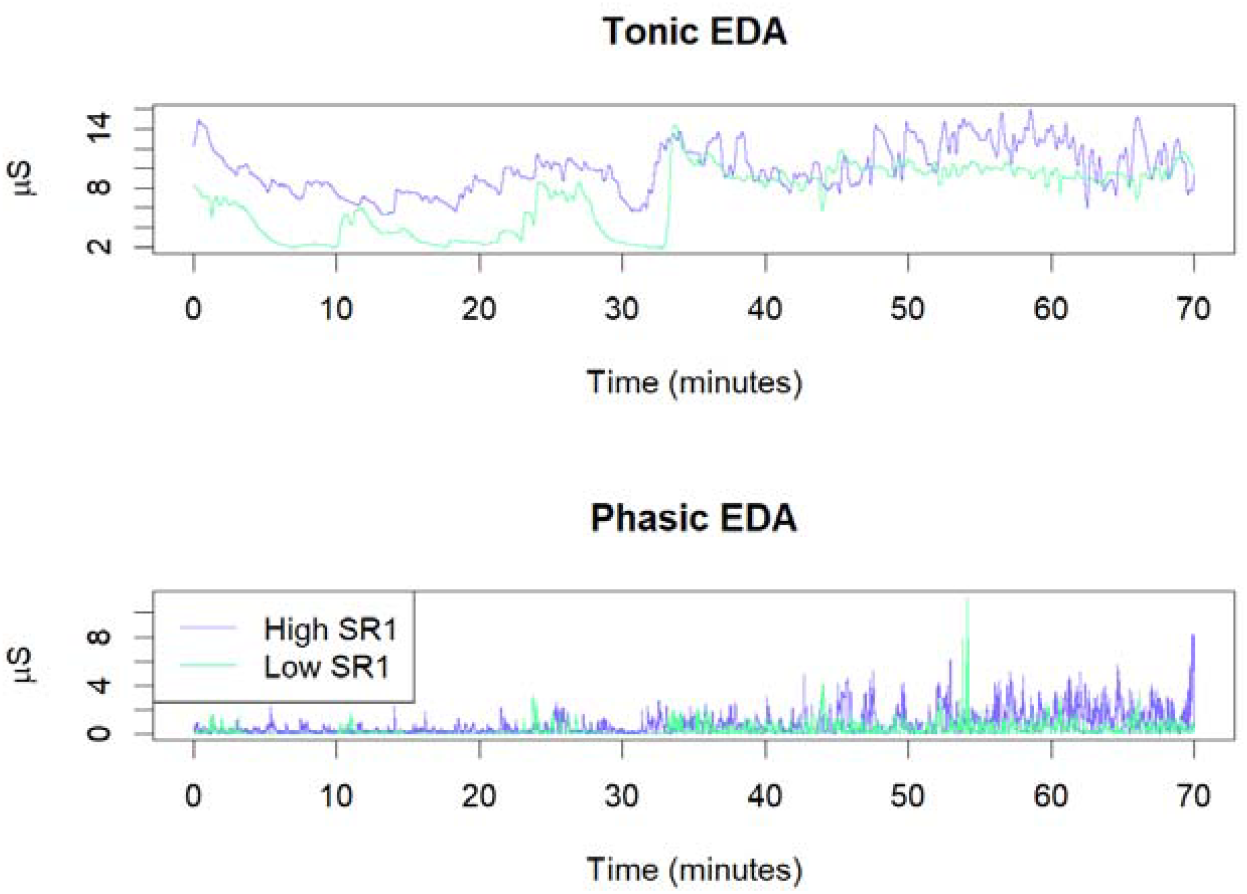
An example of tonic and phasic EDA responses for one high sensory responsivity (SR1) student versus one low sensory responsivity (SR1) student in the classroom study.

To assess whether sensory responsivity predicted tonic or phasic EDA levels in a higher sensory input environment (i.e., a classroom context) compared to the quiet lab, a linear mixed-effects model was estimated with tonic and phasic EDA as outcomes.

This model controlled for spatial ability, gender, EDA type (tonic or phasic), sensory level (high or low), an interaction between the EDA type and sensory level, an interaction between sensory level and the SR factor, and another interaction between the EDA type and the SR factor, and investigated main effects from the SR factor (two different models, as these variables demonstrated high multicollinearity in the classroom context as well (r = .87). Age was not included in this model as all students were within one year of each other in age as they were all in year 12 in upper secondary education.

For the SR1 model, spatial ability, gender, and sensory level were not significant predictors of EDA, and neither were the interactions between sensory level and the SR factor, nor the interaction between sensory level and EDA type. For the SR2 model, there were no significant predictors except the EDA type (b = -6.391, SE = .342, p < .001), thus only the results from the SR1 model are reported further.

A likelihood ratio test comparing the full mixed model for SR1 and a reduced model without the non-significant variables revealed that the full model (χ^2^(5) = 2.47, p = .650) did not provide a significantly better fit to the data than the reduced model without the additional predictors. This comparison therefore justified dropping the spatial ability, gender, sensory level, the interaction between sensory level and the SR factor, and the interaction between sensory level and EDA type and proceeding with the simpler model.

The simple mixed-effects models demonstrated significant contributions to the EDA outcome from all variables. The conditional R^2^ was .748, while the marginal R^2^ was .692. Model diagnostics indicated no violations of linearity, homoscedasticity, or normality of residuals. The variance inflation factors were all below 2, suggesting that multicollinearity did not bias the estimates.

The linear mixed effects model for SR1 revealed significant main effects of SR1 group (b = 1.747, SE = .566, p = .003), EDA type (b = -5.615, SE = .465, p < .001). The model also identified significant interaction effects between SR1 group and EDA type (b = -1.599, SE = .667, p < .018). The results indicate that only SR1 and EDA type significantly influenced EDA responses, with significant interactions between these variables.

Post-hoc pairwise comparisons with Bonferroni-Holm correction revealed that the difference between the SR1 groups was significant only for tonic EDA. Specifically, for tonic EDA, the high SR1 group showed significantly higher EDA responses than the low SR1 group (estimate = -1.747, SE = .579, t(79.2) = -3.017, p = .003). In contrast, no significant differences were found between SR1 groups for phasic EDA (estimate = -.148, SE = .579, t(79.2) = -.256, p = .797). These results indicate that the SR1 group differences are specific to tonic EDA responses, but in the opposite direction from in a quiet lab environment, as shown in Figure 11.

**Figure 11.**
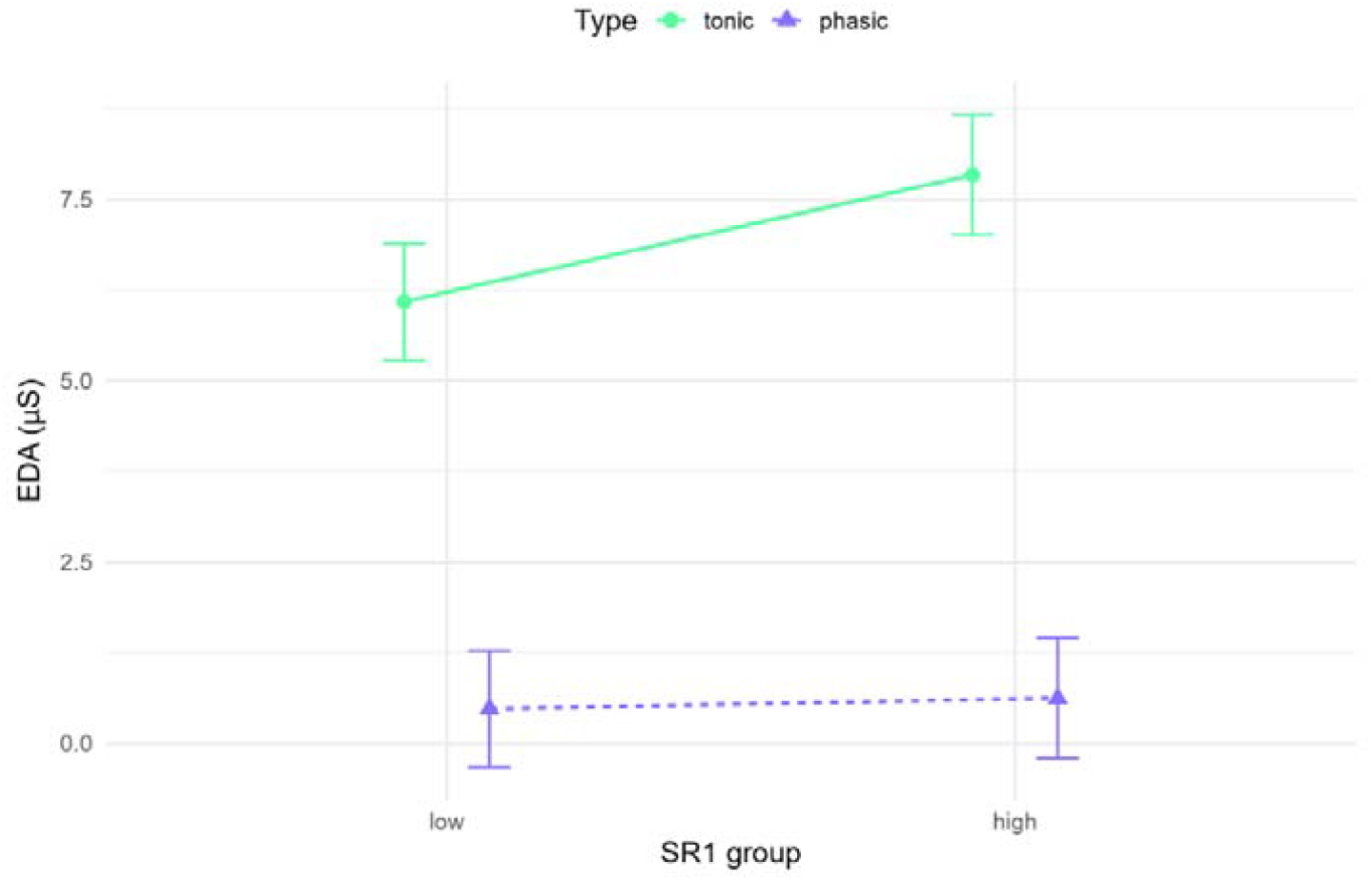
Tonic and phasic EDA response in the classroom study for the SR1 median split groups.

## 4. Discussion

The discussion is structured according to the two research hypotheses, by first focusing on whether the highest SR groups have lower levels of sensory gating in section 4.1, then focusing on whether the highest SR groups have higher levels of sympathetic activation in section 4.2, before discussing implications and future research directions based on the findings in section 4.3.

### 4.1 H1: The highest SR groups have lower levels of sensory gating

This hypothesis was not supported by the results, as the high SR groups demonstrated higher levels of sensory gating than the low SR groups. Because sensory gating worsens with higher stress levels (Johnson & Adler, 1993; Xin et al., 2021) and the high SR groups had the lowest sympathetic activation in the lab (i.e., a measure of stress), the current results are in line with previous research.

Moreover, because sensory gating was increased for the high SR groups, and sensory gating has been linked to attention (Golubic et al., 2019; Jones et al., 2016; Wan et al., 2008), these results may indicate that the high SR students demonstrate improved attention in the low sensory input condition. However, modest effect sizes limit the practical significance of this finding for the general student population. Nevertheless, students with very high sensory responsivity may substantially benefit from less sensory intensive environments, which could enhance their ability to attend to relevant stimuli.

### 4.2 H2: The highest SR groups have higher levels of sympathetic activation, and the difference depends on the sensory intensity of the environment

This hypothesis was partly supported by the results, as only tonic EDA levels differed based on sensory responsivity levels, and that the direction of the effect differed based on the sensory intensity of the environment.

Both tonic and phasic EDA levels assess sympathetic activation (Boucsein, 2012), where tonic EDA reflects the slow-varying baseline signal (Villanueva et al., 2018) and phasic EDA is considered a more immediate and reactive response and has been linked to cognitive load (Pijeira-Díaz et al., 2018). There were no significant differences in phasic sympathetic activation in the lab or in the classroom study based on sensory responsivity, which indicates that cognitive load levels were similar for students with high and low sensory responsivity.

However, the lab study revealed that high sensory responsivity groups demonstrated lower tonic sympathetic activation across all activities. Conversely, the classroom study showed elevated sympathetic activation in the high SR1 group. This discrepancy suggests that while high SR students maintain a relaxed baseline in controlled environments, they experience heightened sympathetic activation in sensory-intensive settings such as classrooms. This physiological response may contribute to unhealthy stress levels and impaired attention, potentially explaining the elevated risk for disengagement and school dropout in this population (Appendix A). Future research should investigate whether sensory input reduction improves retention and engagement among high SR students.

### 4.3 Implications and further research

Overall, these results are in line with the theory of differential susceptibility, where it has been found that those most susceptible to poor outcomes in an unsupportive environment also are most susceptible to good outcomes in supportive environments, i.e., a plasticity approach (Belsky & Pluess, 2009; Ellis et al., 2011). There is genetic meta-analytical empirical support of differential susceptibility (Bakermans-Kranenburg & van IJzendoorn, 2015; van IJzendoorn & Bakermans-Kranenburg, 2015). Calls for further research into its neural mechanisms (Homberg & Jagiellowicz, 2022) and the development of robust measures and examination of underlying mechanisms to inform therapeutic interventions (Assary et al., 2023) have been extended.

The SR scale can contribute to identify individuals’ sympathetic activation based on sensory responsivity, where those high in the trait are more sympathetically reactive to stimuli (i.e., lower in the lowest sensory input conditions and highest in the highest sensory input conditions). This is in line with research on sympathetic activation based on differential susceptibility, where the same pattern was identified (Boyce, 2016). Thus, sensory responsivity may be a candidate for further differential susceptibility research while concurrently providing a suggested mechanism for both positive and negative outcomes through the mediation of stress levels. However, replication of the current results should be sought for wider populations and in a greater variety of contexts.

The full EDA signal (i.e., the sum of the tonic and phasic components) is used in physiological stress research and is mainly affected by the tonic EDA component, as this is typically one order of magnitude larger than the phasic component (Boucsein, 2012). Elevated stress levels have been found to increase risk of multiple conditions, such as coronary artery disease (Boieriu et al., 2024; Yang et al., 2015), atopic dermatitis and psoriasis (Hall et al., 2012), and several autoimmune diseases (Sharif et al., 2018), through the activation of the sympathetic nervous system. Because people high in sensory responsivity demonstrate higher levels of tonic sympathetic activation in all situations but those in a very quiet lab environment, the current study implies that high sensory responsivity may be a risk factor for developing these conditions, in particular if they are exposed to many loud/sensory intensive environments. There is also an increased prevalence of autoimmune conditions for people with ADHD (Chen et al., 2017; Hegvik et al., 2022) and ASD (Chen et al., 2013; Villarreal et al., 2024; Zerbo et al., 2015). This suggests a potential link between sensory responsivity and increased stress levels, as the two former groups are known to have sensory issues, which should be investigated in future studies. This may entail including sensory responsivity assessment in stress-related research because of its moderating role in sympathetic activation.

## Conclusions

This study introduced the concept of sensory responsivity as measured by the sensory responsivity scale. The results showed that tonic sympathetic activation was elevated for high SR groups when sensory intensity was high/normal. Conversely, in very low sensory intensity environments, high SR students demonstrated reduced sympathetic activation and improved sensory gating compared to low SR peers. This indicates improved attention and lower stress levels for high SR students in very low sensory intensive environments compared to low SR students. However, for higher sensory intensive environments such as classrooms, high SR students demonstrated increased stress levels compared to low SR students. Such increased stress levels can increase disease risk for and may explain increased disengagement and school absence for high SR students.

