## Appendix A for "Sensory responsivity and its connection to sympathetic activation and deactivation: implications for stress and attention"

The sensory responsivity (SR) scale development follows a slightly adapted variant of the methodology by Author and colleague (2025) using multidimensional item response theory (IRT) as reported below.

**1. Materials and methods**

**1.1 SR scale design**

36 qualitative items describing sensory responsivity based on literature describing related traits such as SPS (Aron & Aron, 1997; Greven et al., 2019), sensory issues in ASD and ADHD (Ben-Sasson et al., 2019; Dellapiazza et al., 2021; Schaaf et al., 2015; Scheerer et al., 2024), and sensory sensitivities in the gifted (Beljan et al., 2006) was developed.

These items consisted of responsivity items based on several sensory systems:

- Visual responsivity
- Auditory responsivity
- Muscular responsivity (movement and proprioception)
- Tactile responsivity
- Gustatory and olfactory responsivity
- Bodily responsivity (interoception)

Two independent experts evaluated the items for theoretical suitability and thus reduced the number of items to 27, still including all the original sensory systems. The students answered the items on a five-point Likert scale, with the categories “Very poor fit”, “Somewhat poor fit”, “Neutral”, “Somewhat good fit”, and “Very good fit”. These categories were initially coded 1, 2, 3, 4, and 5, respectively. See Table 4 for an overview of the items.

**1.2 Other measures**

The HSP scale (Aron & Aron, 1997) consists of 27 items with a seven-point Likert scale. It has been validated on Norwegian populations (Listou Grimen & Diseth, 2016; Trå et al., 2022) and is described in the main document as follows:

“This scale includes items on positive or negative, emotional or cognitive responses to stimuli such as art, loud noises, smells, and fabrics (Greven et al., 2019). Examples of items are: ‘Do other people's moods affect you?’, ‘Do you make a point to avoid violent movies and TV shows?’, and ‘Are you conscientious?’”

The Negative School Engagement Behaviour (NSEB) subscale aims to identify students' negative engagement behaviour in school (Skinner et al., 2009). Negative school engagement, or school disengagement, is a risk factor for school dropout (Author & colleagues, 2024). The teacher report version of the NSEB scale has been validated for use in a Swedish school context (Ritosa et al., 2020), which is similar to the Norwegian context and language in the current study. The NSEB scale consists of five items that are scored on a four-point Likert scale.

The school absence level was self-reported by the students at five levels.

**1.3 Participants**

The first sample comprised 293 students aged 9 to 16 (grades 5 to 10) from a semi-rural municipality with a small city with a population of around 20,000 in southwest Norway. The second sample included 242 students from the same age group, from a suburban municipality in southeast Norway with a population of approximately 40,000. Table 1 provides an overview of both samples. Students were recruited through their municipalities, with participation from entire classes. The samples were combined and then randomly divided into two groups to facilitate exploratory and confirmatory model estimation.

**Table 1.** The two student samples by self-reported gender and grade.

| Number of participants | Male | Female | Unidentified | Grade 5 |  | Grade 6 | | | Grade 7 | Grade 8 | Grade 9 | Grade 10 |
| --- | --- | --- | --- | --- | --- | --- | --- | --- | --- | --- | --- | --- |
| 293 | 145 | 131 | 17 | 63 | | | 55 |  | 26 | 71 | 47 | 31 |
| 242 | 113 | 113 | 16 | 22 | | | 20 |  | 19 | 70 | 109 | 2 |

The included age range covers a period of much cognitive development, e.g., the reading level changes during this period and it is important to incorporate this in scale development (Gorsuch & Venable, 1983). To accommodate this variability, items were designed to be concise and accessible for younger participants while remaining relevant for older participants.

**1.4 Data collection**

Data was collected through an electronic form of the SR scale (i.e., Nettskjema), and all items were mandatory. Thus, there was no missing data in the data set.

**1.5 Data analysis**

I used R version 4.3.1 (R Core Team, 2023) with R studio version 2023.12.1 for the statistical analysis and the R package mirt (Chalmers, 2012) to estimate the IRT models.

**1.5.1 Descriptive statistics**

Item-to-total score Pearson correlations and item mean scores were used as an initial screening of item quality. If any items had an item-to-total score Pearson correlation lower than 0.3, they were selected for further review. Then Cronbach’s alpha was estimated to obtain a lower-bound estimate of the reliability of the sum scores.

**1.5.2 Dimensionality assessment**

Dimensionality was evaluated by randomly splitting the data set into two equal-sized samples: one sample for the exploratory dimensionality analysis and the other sample for the confirmatory analysis based on the results from the exploratory analysis. Splitting the data resulted in sample A with 268 respondents and sample B with 267 respondents. Initial analysis revealed that the three middle categories were often overlapping, and I thus combined the middle three categories into one. I performed a full-information exploratory IRT analysis with the recoded data using sample A by fitting unrestricted IRT models with one - five dimensions and selecting the model with the lowest Bayesian Information Criteria (BIC) (Cho et al., 2016; Kim et al., 2019) for confirmatory analysis.

**1.5.3 IRT estimation and evaluation**

IRT models, like confirmatory factor analysis, treat constructs such as SR level as latent variables that are not directly observable. These models assume that the probability of a participant endorsing an item depends on two parameter sets: the individual’s position on the SR level continuum (person parameter) and the characteristics of each item (item parameters).

Following the exploratory IRT analysis, I fitted a confirmatory IRT model with sample B and assessed both model and item fit. Model fit was evaluated using several criteria: the M2 hypothesis test (Liu et al., 2016), root mean square error of approximation (RMSEA), and standardized root mean squared residual (SRMSR) statistics. I considered a significance level of 0.05 for the M2 test of absolute fit and used RMSEA < 0.06 and SRMSR < 0.08 as thresholds for good approximate fit (Cai & Monroe, 2014; Maydeu-Olivares & Joe, 2006). Item fit was assessed using Sχ^2^ hypothesis tests (Kang & Chen, 2008) with a Bonferroni-adjusted significance level of 0.05.

**1.5.4 Psychometric properties of the SR scale**

I evaluated the measurement properties of the scale using the estimated IRT model and estimated the reliability of the IRT ability scores with model-based reliability coefficients (Cheng et al., 2012).

**1.5.5 Correlations with external variables**

To obtain a clear interpretation of the SR scale scores, we estimated correlations between the latent ability scores of the SR scale and the HSP scale, between the latent ability scores of the SR scale and the sum scores of the NSEB scale, and between the latent ability scores of the SR scale and the school absence level.

**2. Results**

None of the 27 items of the SR scale had a point biserial item-total correlation below 0.3, so no items were removed from further analysis. I used an exploratory dimensionality analysis where a fully unrestricted graded response unidimensional IRT model was estimated. To investigate dimensionality, graded response models with two - five dimensions were also estimated and compared to the unidimensional model. According to the BIC, the best model was the two-dimensional model (BIC = 10704) compared to the unidimensional model (BIC = 10741), the three-dimensional model (BIC = 10714), the four-dimensional model (BIC = 10794), and the five-dimensional model (BIC = 10869).

I then inspected visual item fit by viewing trace plots to evaluate if categories should be collapsed to better fit the data. The plots demonstrated substantial overlaps between the middle three categories. Therefore, it was decided to collapse the middle three categories into only one, as there was no theoretical reason to keep all five categories. The first category was named “Very poor fit” and was coded as 0, the three combined categories were named “Somewhat poor fit”, “Neutral”, and “Somewhat good fit” and were coded as 1, whilst the remaining category was called “Very good fit”, and was coded as 2.

The confirmatory model evaluation also collapsed the middle three categories into one before the computation of the item mean, M = 28.2, standard deviation, SD = 7.69, and Cronbach’s α = 0.89. Item mean and item-total score correlations are given in Table 2.

**Table 2.** The item means, item-total score correlation (pBIS), and internal consistency (α) if the item is deleted.

| Item | Mean | pBIS | α |
| --- | --- | --- | --- |
| 1 | 0.96 | 0.51 | 0.89 |
| 2 | 1.19 | 0.43 | 0.89 |
| 3 | 1.01 | 0.45 | 0.89 |
| 4 | 1.00 | 0.58 | 0.89 |
| 5 | 1.03 | 0.46 | 0.89 |
| 6 | 1.16 | 0.34 | 0.89 |
| 7 | 0.84 | 0.50 | 0.89 |
| 8 | 0.99 | 0.49 | 0.89 |
| 9 | 0.92 | 0.53 | 0.89 |

| Item | Mean | pBIS | α |
| --- | --- | --- | --- |
| 10 | 0.91 | 0.52 | 0.89 |
| 11 | 0.91 | 0.52 | 0.89 |
| 12 | 1.16 | 0.54 | 0.89 |
| 13 | 1.12 | 0.50 | 0.89 |
| 14 | 1.03 | 0.50 | 0.89 |
| 15 | 1.24 | 0.40 | 0.89 |
| 16 | 0.95 | 0.37 | 0.89 |
| 17 | 1.11 | 0.53 | 0.89 |
| 18 | 1.19 | 0.32 | 0.89 |

| Item | Mean | pBIS | α |
| --- | --- | --- | --- |
| 19 | 1.18 | 0.32 | 0.89 |
| 20 | 1.02 | 0.54 | 0.89 |
| 21 | 0.94 | 0.42 | 0.89 |
| 22 | 1.06 | 0.52 | 0.89 |
| 23 | 1.14 | 0.41 | 0.89 |
| 24 | 1.05 | 0.45 | 0.89 |
| 25 | 1.04 | 0.50 | 0.89 |
| 26 | 1.17 | 0.41 | 0.89 |
| 27 | 0.91 | 0.41 | 0.89 |

Then, a model evaluation based on the model from the first sample with a confirmatory analysis was performed. The two-dimensional model from the exploratory analysis was applied to the second dataset from the combined sample, sample B. I assessed model-data fit by the M2-statistic, which provided the values RMSEA = .06 (90 % CI = [.05, .07]), SRMR = .07, demonstrating good fit (Maydeu-Olivares, 2013). The reliability of the expected a posteriori (EAP) ability estimates (Bock & Mislevy, 1982) resulted in acceptable values for both factors, namely, SR1 = 0.81 and SR2 = 0.77 (Maydeu-Olivares, 2013).

The S-χ^2^ item fit test was performed to evaluate the fit of the individual items with the Bonferroni corrected value of the probability, p = .0019 (i.e., a confidence level of 5 %). As this value is conservative, I considered any items with p-values p > .002 to demonstrate adequate fit. Table 3 shows that all items demonstrated adequate fit.

**Table 3.** The results of the S-χ^2^ test of item fit.

| Item | S-$\chi^{2}$ | Probability |
| --- | --- | --- |
| 1 | 17.2 | 0.44 |
| 2 | 22.6 | 0.16 |
| 3 | 19.8 | 0.34 |
| 4 | 18.3 | 0.43 |
| 5 | 22.5 | 0.26 |
| 6 | 15.1 | 0.52 |
| 7 | 37.5 | 0.03 |
| 8 | 24.4 | 0.14 |
| 9 | 21.9 | 0.19 |

| Item | S-$\chi^{2}$ | Probability |
| --- | --- | --- |
| 10 | 28.1 | 0.04 |
| 11 | 19.8 | 0.23 |
| 12 | 16.8 | 0.33 |
| 13 | 22.1 | 0.18 |
| 14 | 33.7 | 0.03 |
| 15 | 23.0 | 0.11 |
| 16 | 31.2 | 0.09 |
| 17 | 15.5 | 0.41 |
| 18 | 25.5 | 0.04 |

| Item | S-$\chi^{2}$ | Probability |
| --- | --- | --- |
| 19 | 29.1 | 0.05 |
| 20 | 9.1 | 0.82 |
| 21 | 31.0 | 0.03 |
| 22 | 26.1 | 0.07 |
| 23 | 19.2 | 0.44 |
| 24 | 16.9 | 0.53 |
| 25 | 23.1 | 0.15 |
| 26 | 23.9 | 0.09 |
| 27 | 40.9 | 0.02 |

I then visually evaluated item fit by inspecting item trace plots to check for non-overlap in categories, and no overlaps were identified. Examples of item trace planes are shown in Figure 1.

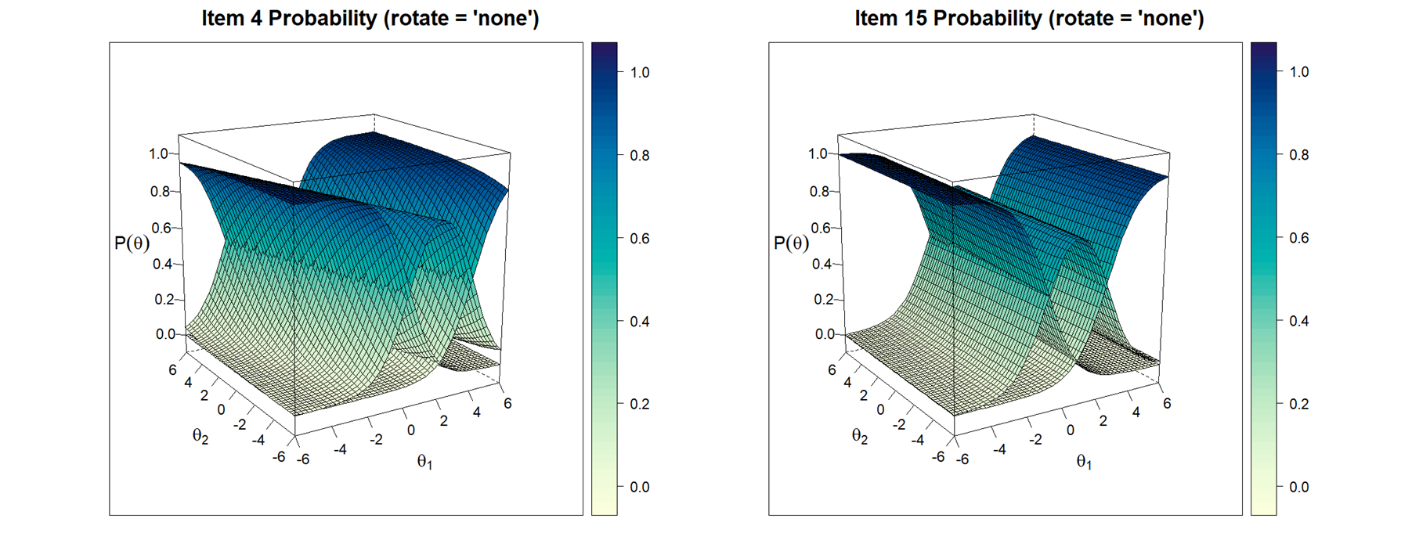

**Figure 1.** Examples of item trace curves are shown for items 4 and 15.

The factor loadings for the two factors, namely SR1 and SR2, were estimated with oblimin rotation, with the corresponding factor correlation = 0.56. As seen in Table 4, the SR1 factor mainly describes exteroception, i.e., the sensory input from the outside world (visual, auditory, gustatory, and olfactory sensory input), whereas the SR2 factor mainly describes interoception, i.e., the sensory input from the body itself (muscular, tactile, and bodily sensory input).

**Table 4.** An overview of the items in the SR scale and their corresponding factor loadings. Loadings over 0.3 are highlighted.

| Item | Factor loadings | |
| --- | --- | --- |
|  | SR1 | SR2 |
| **Visual responsivity** |  |  |
| 1. It is very important to me that it is neither too bright nor too dark in a room. | **0.54** | 0.22 |
| 2. Flashing lights are very annoying. | **0.38** | 0.21 |
| 3. It is very uncomfortable to be surrounded by clutter. | **0.54** | 0.05 |
| 4. When I enter a room, I immediately notice bright lights. | **0.52** | 0.12 |
| **Auditory responsivity** |  |  |
| 5. It often becomes too noisy in the classroom for me. | **0.41** | **0.31** |
| 6. I am good at hearing the quietest of sounds. | **0.32** | 0.25 |
| 7. I often think that the movie theatre sound is too loud. | **0.69** | -0.17 |
| 8. It is very uncomfortable to hear sirens close by. | **0.69** | -0.08 |
| 9. I think that some people breathe noisily. | **0.62** | 0.14 |
| **Muscular responsivity (movement and proprioception)** |  |  |
| 10. I prefer to move carefully. | **0.61** | 0.16 |
| 11. I notice more easily than others that my body feels tired. | **0.28** | **0.46** |
| 12. Having a comfortable place to sit is important to me. | 0.13 | **0.57** |
| 13. It is unpleasant to stand for a long time. | 0.04 | **0.64** |
| 14. I quickly notice when my muscles get tired. | 0.09 | **0.57** |
| **Tactile responsivity** |  |  |
| 15. Labels in clothes are often annoying. | **0.32** | **0.31** |
| 16. I’m so ticklish that it almost hurts when somebody tickles me. | **0.47** | 0.11 |
| 17. There are types of clothes or fabrics that feel unpleasant to wear. | 0.25 | **0.60** |
| 18. I only wear comfortable clothes. | 0.08 | **0.59** |
| 19. My shoes must be very comfortable. | -0.27 | **0.73** |
| **Gustatory and olfactory responsivity** |  |  |
| 20. I often notice smells that other people don’t. | **0.47** | **0.44** |
| 21. I can’t eat bitter foods, such as grapefruit and Brussels sprouts. | **0.68** | -0.15 |
| 22. It is difficult for me when others eat food with an unpleasant smell. | **0.52** | -0.04 |
| 23. There are many foods I don’t like to eat. | **0.34** | 0.18 |
| 24. I notice strange tastes more often than others. | **0.56** | 0.14 |
| **Bodily responsivity** |  |  |
| 25. I react strongly to being hungry. | 0.00 | **0.53** |
| 26. I function much more poorly than usual when I am tired. | 0.21 | **0.39** |
| 27. I am more often cold than other people. | 0.24 | 0.22 |

The HSP-scale (Aron & Aron, 1997) for sensory processing sensitivity estimation produced a mean value of M = 109 out of 189 possible points, a standard deviation of SD = 26, and internal consistency of α = 0.91. A three-dimensional graded response model was estimated and assessed by the M2-statistic, which provided the values RMSEA = 0.05 (CI = [0.04, 0.07]), SRMSR = 0.06, demonstrating good fit (Maydeu-Olivares, 2013). Due to non-normal distributions with ties the Kendall correlation of the latent variables (EAP) was estimated and resulted in the correlations listed in Table 5.

**Table 5.** The pairwise correlations between the two factors of the SR scale and the three factors of the HSP scale.

| SR factor | HSP factor | Kendall’s tau | p-value |
| --- | --- | --- | --- |
| SR1 | HSP1 | 0.33 | 7.3e-14*** |
| SR1 | HSP2 | 0.10 | 0.025* |
| SR1 | HSP3 | 0.30 | 6.7e-12*** |
| SR2 | HSP1 | 0.28 | 2.1e-10*** |
| SR2 | HSP2 | 0.13 | 0.0032** |
| SR2 | HSP3 | 0.29 | 1.7e-11*** |

* denotes p-values < 0.05, ** denotes p-values < 0.01, and *** denotes p-values < 0.001.

The negative school behaviour engagement scale has a 4-point Likert-type scale ranging from 1 (not at all true) to 4 (very true) (Skinner et al., 2009). It produced a mean value of M = 10.8 out of 20 possible points, SD = 3.3, and Cronbach’s α = 0.68. No suitable measurement models were identified by either IRT or factor analysis, causing the use of latent scores (EAP estimated) for the STEM scale and sum scores for the engagement scale for correlation check. The Kendall correlation was used due to non-normal ability distributions with ties, which resulted in the values of τ = 0.17 for SR1 (p = 2.1e-08) and τ = 0.17 for SR2 (p = 4.5e-08).

School absence was self-reported by the students at five levels, namely, ‘nothing’, ‘little’, ‘ordinary’, ‘larger than most’, and ‘very large’. These levels were coded as 0, 1, 2, 3, and 4, respectively. The Kendall correlation was used due to non-normal distributions with ties, which resulted in the values of τ = 0.09 for SR1 (p = 0.0058) and τ = 0.07 for SR2 (p = 0.025).
